# Extraction-Free Whole Transcriptome Gene Expression Analysis of FFPE Sections and Histology-Directed Subareas of Tissue

**DOI:** 10.1101/412015

**Authors:** Christy L Trejo, Miloš Babić, Elliot Imler, Migdalia Gonzalez, Sergei Bibikov, Peter J Shepard, Harper C VanSteenhouse, Joanne M Yeakley, Bruce E Seligmann

**Affiliations:** BioSpyder Technologies, Incorporated, Carlsbad, California, United States of America

## Abstract

We describe the use of a ligation-based targeted whole transcriptome expression profiling assay, TempO-Seq™, to profile formalin-fixed paraffin-embedded (FFPE) tissue, including H&E stained FFPE tissue, by directly lysing tissue scraped from slides without extracting RNA or converting the RNA to cDNA. The correlation of measured gene expression changes in unfixed and fixed samples using blocks prepared from a pellet of a single cell type was R^2^ = 0.97, demonstrating that no significant artifacts were introduced by fixation. Fixed and fresh samples prepared in an equivalent manner produced comparable sequencing depth results (+/-20%), with similar %CV (11.5 and 12.7%, respectively), indicating no significant loss of measurable RNA due to fixation. The sensitivity of the TempO-Seq assay was the same whether the tissue section was fixed or not. The assay performance was equivalent for human, mouse, or rat whole transcriptome. The results from 10 mm^2^ and 2 mm^2^ areas of tissue obtained from 5 μm thick sections were equivalent, thus demonstrating high sensitivity and ability to profile focal areas of histology within a section. Replicate reproducibility of separate areas of tissue ranged from R^2^ = 0.83 (lung) to 0.96 (liver) depending on the tissue type, with an average correlation of R^2^ = 0.90 across nine tissue types. The average %CVs were 16.8% for genes expressed at greater than 200 counts, and 20.3% for genes greater than 50 counts. Tissue specific differences in gene expression were identified and agreed with the literature. There was negligible impact on assay performance using FFPE tissues that had been archived for up to 30 years. Similarly, there was negligible impact of H&E staining, facilitating accurate visualization for scraping and assay of small focal areas of specific histology within a section.

## Introduction

Gene expression profiling of tissue is vitally important for understanding both normal and disease processes. Tissue can be prepared as snap frozen blocks or prepared as formalin fixed paraffin embedded (FFPE) tissue blocks, then sectioned and assayed. Frozen tissue blocks are amenable to gene expression assays, but not without significant problems. The samples are difficult to handle and transport, as they must be kept frozen from the moment of collection onwards. Due to difficulties with staining and archiving, the general pathological practice is to collect FFPE blocks rather than freeze (1, 2).

Assays of gene expression in formalin-fixed paraffin-embedded tissues have historically been complex and problematic, and have often provided subpar data (1-5). Extraction of RNA from FFPE is typically low yield compared to fresh or frozen tissue, and the resulting RNA is highly fragmented and degraded, leading to poor performance in the usual methodologies which rely on reverse-transcription of RNA. The problem is particularly significant for archival FFPE samples, where RNA degradation is generally more pronounced. Vast collections of such samples are currently present in various hospitals and research centers around the world, and these samples are often matched with detailed clinical and outcome data. Similar archives exist for animal tissues from experimental studies performed over many decades. Yet, the treasure trove of gene expression data available in such archives has largely remained out of reach.

Additional problems are caused by tissue heterogeneity, as samples usually contain many different cell types and associated histology within a section, where the cell type or histology of interest may represent a small percentage of the total. Current FFPE RNA extraction methods typically require multiple complete sections of tissue to be processed together to recover sufficient material for transcription assays (6). Therefore, a method that does not require the extraction of RNA, is not sensitive to fragmentation, and which can be used to profile small focal histologically and distinct areas of archived (not just fresh) FFPE has tremendous potential to advance science.

Two other commercial platforms enable investigators to profile FFPE without extraction of RNA, but both require dedicated hardware and neither permit the whole transcriptome to be profiled. nCounter™ (NanoString, Inc.) is limited to profiling the expression of up to 700 genes. EdgeSeq™ (HTG Molecular, Inc.) is limited to a few thousand genes. Thus, to use either of these methods the investigator must already know the set of genes to monitor in a focused assay, which means they have to first carry out a method such as RNAseq (or microarray) which necessitates extracting RNA from FFPE, or using fresh or frozen tissue. Translating such data to a different platform is typically problematic. Hence, a method that enables profiling of the whole transcriptome, some 20,000 genes, from FFPE without RNA extraction, without use of dedicated hardware, and then (as desired) selecting genes to formulate a focused assay on the same platform, would be a significant advance.

In this study, we extend the use of the targeted, ligation-based Templated Oligo Sequencing (TempO-Seq™) whole transcriptome assay to FFPE samples (7). The experiments described were carried out using human, mouse, and rat whole transcriptome panels. Since it relies on probe hybridization rather than reverse transcription, TempO-Seq chemistry is highly resistant to RNA fragmentation and degradation, making it perfectly suited for fixed tissue samples. We show that FFPE samples can produce gene expression data on par with fresh tissues and cell cultures, that decades-old archival samples can be successfully processed without laborious extraction and purification methods, and that such tissues can even be H&E stained so that small focal areas of interest can be profiled independently of the surrounding tissue.

## Materials

TempO-Seq FFPE assay reagents are commercially available from BioSpyder Technologies, Inc. The components of the kit are proprietary and consist of 2X Lysis buffer, FFPE Protease, FFPE nuclease, species-specific detector oligo pools designed to recognize the whole transcriptome, and buffers necessary for annealing, nuclease clean up, ligation, amplification, and library generation. For library purification, we used the NucleoSpin Gel and PCR Clean-up kit (Macherey-Nagel cat # 740609.50). Molecular biology grade light mineral oil was sourced from Sigma. Phosphate Buffered Saline (PBS), Ca^2+^ and Mg^2+^ free, was purchased from Thomas Scientific. Molecular biology grade water and TE were purchased from Invitrogen. Ethanol was sourced from Decon Laboratories. Neutral buffered formalin was purchased from VWR (16004-128). All reagents, tips, plates, and reservoirs were RNase free.

## Methods

### Hematoxylin and Eosin staining

Slides were deparaffinized using the Leica Bond Dewax solution by soaking in a Coplin staining jar for three minutes. Slides were washed 3 times in 100% ethanol, then either air dried, or continued through H&E staining with the following protocol: rehydrated in distilled water for three minutes; immersed in hematoxylin solution (Leica hematoxylin 560 diluted 1:6 in distilled water) for three minutes; washed three times in distilled water; immersed briefly in 0.1x PBS; washed three more times in distilled water; soaked in 70% Ethanol for two minutes; immersed in Alcoholic Eosin Y with Phloxine (Sigma HT110332) for three minutes; washed in 100% ethanol three times then air dried.

### Cell lysates and FFPE pellets

MCF7 and MDA-MB-231 cells were obtained from ATCC and grown in RPMI supplemented with 10% FBS. Fresh lysates were prepared by washing cells with 1X Ca^2+^ and Mg^2+^ free PBS, then lysing in 1X TempO-Seq lysis buffer in PBS at 2,000 cells per µL. Lysates were incubated for 10 minutes at room temperature, followed by storage at −80°C. For FFPE cell pellets, live cells were washed twice with 1X PBS, and then fixed with 1% formaldehyde in 1X PBS for 30 minutes at room temperature. Cells were then embedded by the TACMASR (University of Arizona Tissue Acquisition and Cellular/Molecular Analysis Shared Resource).

### FFPE tissues

All human tissue was sourced from the University of Arizona Cancer Center Biorepository. Prostate samples were sourced from the UACC Prostate Biorepository. Human samples were consented for research use after clinical testing and deidentified before receipt. Archival samples were stored as FFPE blocks from the year indicated until 2016, at which point a pathologist identified homogenous tumor regions. 5 µm thick sections were then cut and mounted, and slides were stored until 2018. For these experiments, 25 mm^2^ areas were cut from serial sections to represent biological replicates.

Mouse tissue was provided as a gift from Kathleen Scully and Pamela Itkin-Ansari of the Sanford Burnham Prebys Medical Discovery Institute (La Jolla, Ca.). All mice were wildtype adults of the C57BL/6 genetic strain. Tissues were fixed in 10% neutral buffered formalin at 4°C for 24 hours, then moved to 70% EtOH for 24 hours before embedding. 5 µm thick tissue sections were cut and mounted on Superfrost Plus Micro slides (VWR). Slides were dried overnight at room temperature before processing for FFPE TempO-Seq.

Rat tissues were obtained from Tissue Acquisition and Cellular/Molecular Analysis Shared Resource at University of Arizona Cancer Center. Rats were euthanized with CO^2^ asphyxiation in accordance with American Veterinary Medical Association (AVMA) guidelines. Tissues were fixed in 10% neutral buffered formalin for 24 hours before being transferred to 70% Ethanol prior to embedding. For fixation time studies, samples were kept in 10% NBF for the specified time before moving to ethanol prior to embedding.

### TempO-Seq Assay

The TempO-Seq assay for FFPE samples relies on the standard TempO-Seq chemistry (7-11). The assay (Figure 1) was modified for FFPE samples as follows: An area of interest on a slide-mounted FFPE section (Figure 1A) is scraped from the slide and deposited directly into BioSpyder 1X FFPE lysis buffer (Figure 1B). The sample is then overlaid with molecular biology grade mineral oil and incubated at 95° for five minutes to dissolve the paraffin. FFPE Protease is added and the sample is incubated at 37°C for 30 minutes.

**Figure 1.**
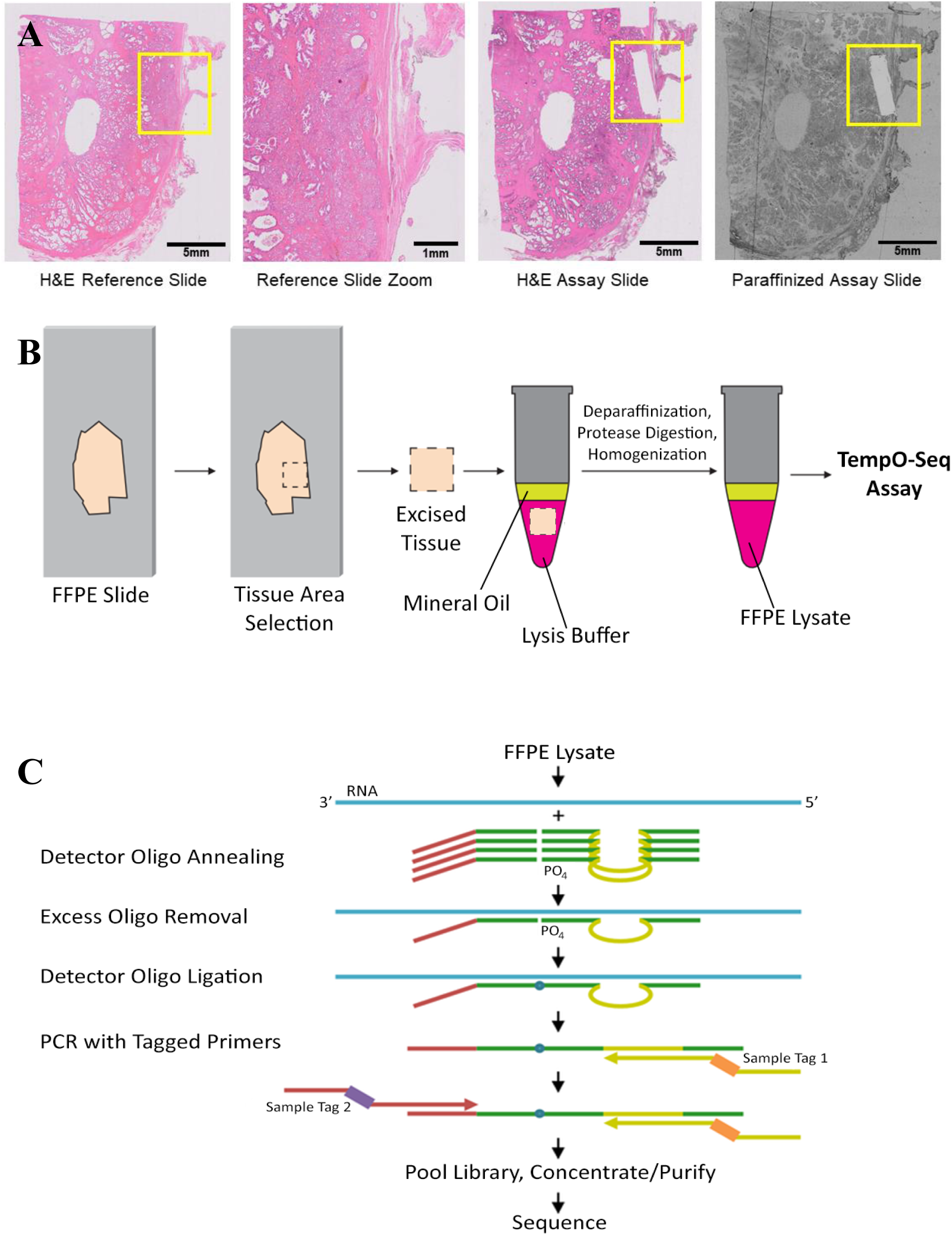
Processing FFPE samples for the TempO-Seq assay. (**A**) Examples of slides processed through FFPE TempO-Seq assay. Left panel shows an H&E stained section, with the yellow box indicating the area of interest. Center left shows an expanded image of this area, demonstrating the mixed histology that would affect the data if the entire area were to be scraped and profiled. Areas identified from a stained tissue section can be scraped from an unstained, paraffinized adjacent section (right), or (if RNase-free reagents are used for staining) directly from a stained section (center right). The scraped areas in this case were ∼ 1 x 5 mm, aligned with the focal histology of interest, and sufficient for gene expression profiling. (**B**) Schematic of the TempO-Seq detector oligo annealing and ligation process. (**C**) An area of interest is manually scraped from mounted FFPE sections. The tissue is added directly into 1X FFPE lysis buffer, overlaid with mineral oil, and then heated at 95°C for 5 minutes. FFPE Protease is added and the sample is incubated and manually homogenized. The processed lysate is then ready for input directly into the annealing step of the TempO-Seq assay.

As depicted in Figure 1B, a 2 µL aliquot of the processed lysate is then added to a microplate well containing a mix of annealing buffer and Detector Oligos (DOs) to measure each targeted gene. DO panels included in this study were designed against the whole transcriptome for human, mouse, and rat (commercially available assays from BioSpyder, Inc.). This mixture was then incubated to anneal probes to the target RNAs. This process is highly resistant to RNA fragmentation (as the DOs anneal to RNA sequences of <100 nt).

Incorrectly bound or unbound DOs are then degraded using FFPE Nuclease, and correctly bound DOs are ligated. After ligation and inactivation of enzymes, the resulting ligated probes are amplified in a PCR step. The PCR primers allow indexing of individual samples, so that hundreds or thousands of samples can be multiplexed within the same sequencing library (Figure 1C).

### Sequencing and Data Analysis

Purified libraries were run on the Illumina NextSeq High 2500 sequencing platform. All data analysis was done using the TempO-SeqR data analysis platform (BioSpyder., Inc.) as follows. After sample demultiplexing using the default Illumina sequencer and bcl2fastq settings, mapped reads were generated by aligning FASTQ files to the ligated DO gene sequences using Bowtie, allowing for up to 2 mismatches in the 50-nucleotide target sequence. For correlation analysis, genes with 20 or more raw counts were log2 transformed and plotted to derive R^2^ values. Differential expression was assessed using the DESeq2 method for differential analysis of count data (12). The count data are first normalized using the DESEq2 function estimateSizeFactor. DESEq2 then computes the probability of differential expression by comparing the relative count level for each condition and the dispersion of the respective counts using a negative binomial model. An adjusted p-value of <0.05 is used as the threshold of significance for differential expression.

The assays used were the human whole transcriptome assay (7) which measures 19,283 genes (21,111 probes); the mouse whole transcriptome assay which measures 23,580 genes (30,147 probes); and the rat whole transcriptome assay which measures 21,119 genes (22,253 probes). Each gene is measured by one or more probes formed by ligation of a DO pair, as previously described (7).

Raw sequencing data in form of FASTQ files, along with aligned gene counts for all samples used in this study are available through GEO (accession number GSE119630).

## Results

### Reproducibility and sample types

To verify assay robustness, we tested tissues from a variety of species and a broad range of tissue types. Data shown here includes FFPE samples of human colorectal cancer, prostate cancer, and pancreatic cancer; rat brain, kidney and liver; and mouse breast, lung, and hindlimb muscle. We chose pancreas tissue due to its relative abundance of endogenous RNases, breast and lung for their low cellularity, and muscle for potential difficulties in digestion. For each sample type, 10mm^2^ areas from 5µm thick slides were lysed, and 10% of the lysate (equivalent to 1 mm^2^ of tissue) was used as input in the whole transcriptome FFPE TempO-Seq assay with species-specific DOs. To gauge assay reproducibility, the same areas of adjacent 5 µm thick sections were independently processed, and gene expression patterns between replicates compared.

Gene expression correlation among biological replicates for all samples types had R^2^ values greater than 0.8, regardless of species. Of the human samples, pancreatic tissue had the highest reproducibility across biological replicates (R^2^ = 0.916), (Figure 2). Human colorectal and prostate cancers had R^2^ values of 0.872 and 0.885, respectively. Mouse breast, lung, and muscle had R^2^ values that exceeded 0.8 (0.891, 0.833 and 0.895, respectively). Rat brain, kidney, and liver all had R^2^ values that exceeded 0.9 (0.926, 0.949, and 0.959), (Figure 2). Across all nine tissue types the average R^2^ was 0.903. Average %CVs for genes with minimum of 10, 50, or 200 counts were 26.7%, 20.3%, and 16.8%, respectively (Table 1).

**Table 1.** Coefficients of variation observed for genes expressing at a minimum level of 10, 50, or 200 counts. Larger variance was observed in samples from tissues known to contain large amounts of RNAses (pancreas), and in tissues with low cellularity and thus low RNA amounts (lung, breast).

|  | >10 | >50 | >200 |
| --- | --- | --- | --- |
| Human Pancreas | 33.44 | 29.80 | 28.88 |
| Human Colon | 27.96 | 22.35 | 19.23 |
| Human Prostate | 27.57 | 19.96 | 15.33 |
| Mouse Breast | 26.45 | 19.67 | 17.01 |
| Mouse Lung | 37.72 | 27.58 | 20.74 |
| Mouse Muscle | 26.06 | 18.79 | 15.18 |
| Rat Brain | 21.33 | 15.96 | 12.98 |
| Rat Kidney | 20.62 | 15.24 | 11.52 |
| Rat Liver | 19.24 | 13.72 | 10.73 |
| Average | 26.71 | 20.34 | 16.84 |

**Figure 2.**
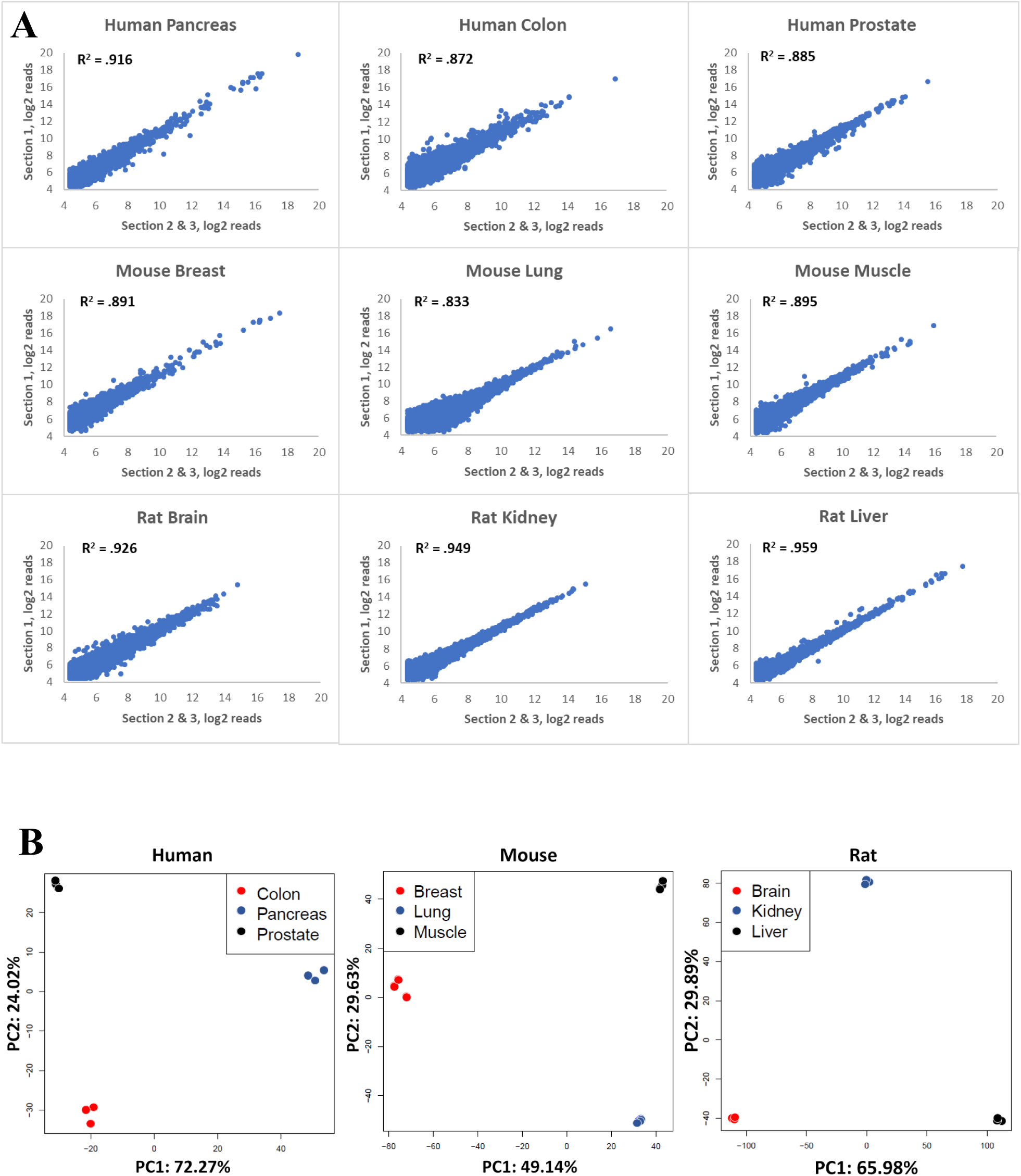
Correlation of gene expression from biological replicates of multiple tissue types. (**A**) FFPE samples of three different tissue types from human, mouse, and rat were used as input into the TempO-Seq FFPE assay. Replicates were obtained by scraping adjacent areas from three different serial sections. R^2^ values were calculated by comparing gene expression of one section to the average of the remaining two. (**B**) Principal component analysis (PCA) of different tissue types of human (colon, pancreas, and prostate), mouse (breast, lung, and muscle), and rat (brain, kidney, and liver) was conducted. The first two principal components account for the majority of variance in the samples and clearly distinguish the different tissue types.

Observed gene expression patterns match the expected transcript abundances. For example, Table 2 shows the highest expressing genes in pancreatic samples and includes expected pancreatic enzymes such as Amylase 2 alpha (AMY2A) and pancreatic lipase (PNLIP), indicating that the amplified and sequenced probes were highly tissue specific. Figure 2B shows a principal component analysis plot (PCA) for these samples, showing clustering of biological replicates and separation of tissue types for each species on both main principal component axes.

**Table 2.** Top 25 highest expressed genes from human pancreatic cancer FFPE samples processed through the FFPE TempO-Seq human whole transcriptome assay.

| Rank | Gene Symbol | Gene Name |
| --- | --- | --- |
| 1 | CTRB1 | chymotrypsinogen B1 |
| 2 | PRSS1 | serine protease 1 |
| 3 | CPA1 | carboxypeptidase A1 |
| 4 | PNLIP | pancreatic lipase |
| 5 | AMY2A | amylase, alpha 2A (pancreatic) |
| 6 | CELA3A | chymotrypsin like elastase family member 3A |
| 7 | AMY2A | amylase, alpha 2A (pancreatic) |
| 8 | CPB1 | carboxypeptidase B1 |
| 9 | CEL | carboxyl ester lipase |
| 10 | CTRB2 | chymotrypsinogen B2 |
| 11 | MTRNR2L12 | MT-RNR2 like 12 |
| 12 | CELA3B | chymotrypsin like elastase family member 3B |
| 13 | CLPS | colipase |
| 14 | PRSS2 | serine protease 2 |
| 15 | CELA2A | chymotrypsin like elastase family member 2A |
| 16 | REG1B | regenerating family member 1 beta |
| 17 | RPL4 | ribosomal protein L4 |
| 18 | AMY1B | amylase, alpha 1B (salivary) |
| 19 | GP2 | glycoprotein 2 |
| 20 | CLPS | colipase |
| 21 | CPA2 | carboxypeptidase A2 |
| 22 | PNLIPRP1 | pancreatic lipase related protein 1 |
| 23 | AMY1A | amylase, alpha 1A (salivary) |
| 24 | SPINK1 | serine peptidase inhibitor, Kazal type 1 |
| 25 | EEF1A1 | eukaryotic translation elongation factor 1 alpha 1 |

### Focal input and sensitivity

Common gene profiling assays generally require RNA purification from large FFPE tissue samples (entire slides or multiple slides). TempO-Seq does not require use of extracted RNA, rather direct sample lysates can be used. Thus, while significant amounts of FFPE are required for RNA extraction, much smaller amounts of FFPE can be assayed as a lysate. The ability to assay very small tissue amounts would spare the use of rare and precious archival FFPE samples, enable profiling of small focal areas with specific pathologies, reduce input for tissues with very low cellularity, and allow profiling of small FFPE samples such as tissue from biopsies or prepared as tissue microarrays. To evaluate the sensitivity and amount of tissue required for TempO-Seq, we tested areas as small as 2 mm^2^ from 5 µm thick tissue sections. The lysis buffer volume was scaled accordingly, so for this input, the amount of tissue in the 2 ul volume that is transferred into the assay was the same as for larger tissue excisions.

We excised both 2 mm^2^ and 10 mm^2^ from 5µm thick mouse liver sections. The correlation of gene expression across biological replicates of the same area excision were similar for 2 mm^2^ and 10 mm^2^ areas, with the R^2^ = 0.969 and 0.95, respectively (Figure 3 A,B). The correlation between 10 mm^2^ and 2 mm^2^ inputs was also very good, with R^2^ of 0.969 (Figure 3 C, average of three samples of each input). These data indicate that the FFPE TempO-Seq assay is highly sensitive and can handle very low input amounts.

**Figure 3.**
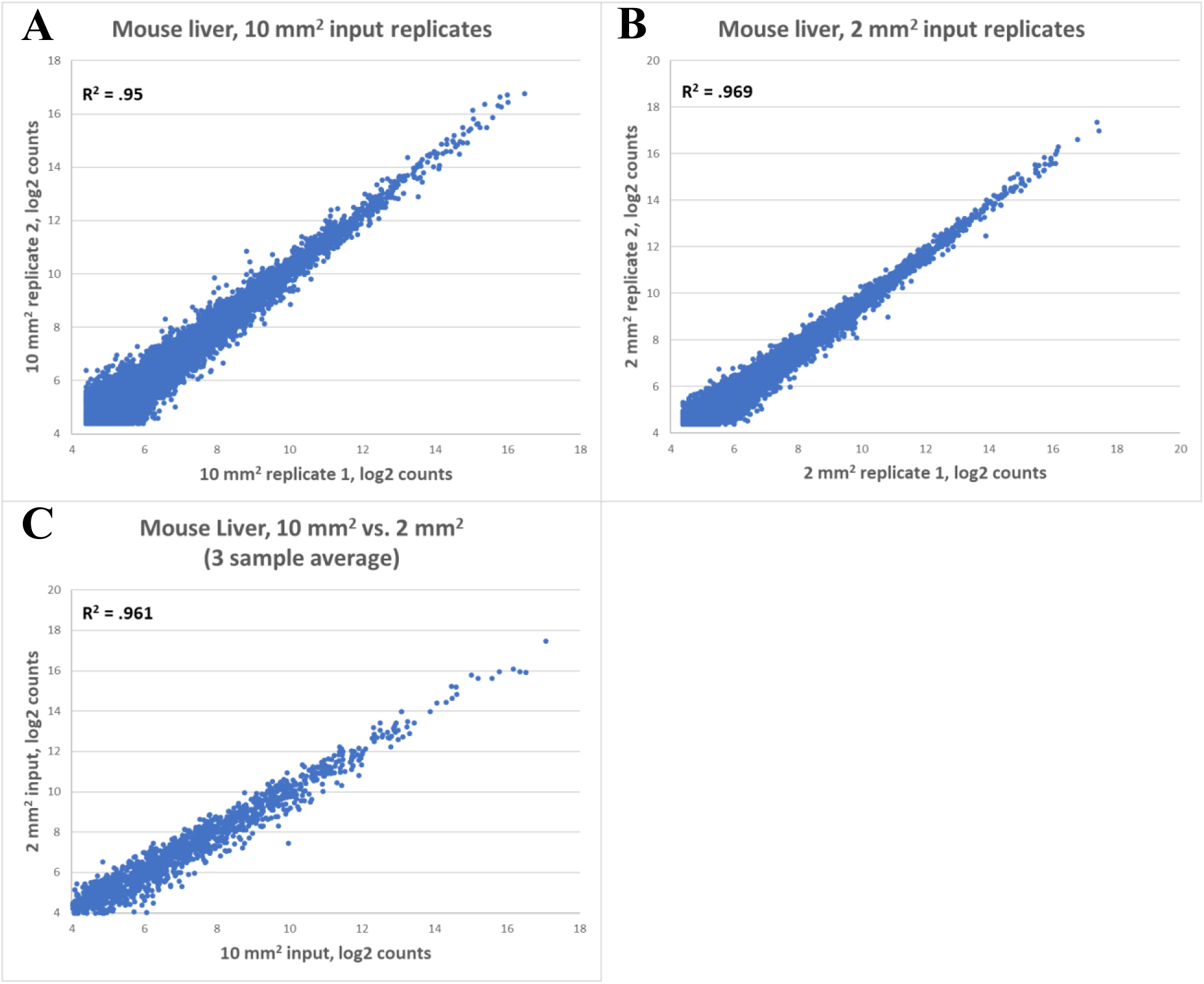
Replicability of 10 mm^2^ and 2 mm^2^ inputs in FFPE TempO-Seq assay. Both (**A**) 10mm^2^ and (**B**) 2mm^2^ areas were scraped from rat liver. Biological replicates were generated by scraping the same area from adjacent tissue and gene expression correlation was calculated. (**C**) Gene expression correlation was calculated between 10 mm^2^ and 2 mm^2^ tissue sections from rat liver.

### Archival tissue

Fixation and paraffin-embedding of tissue allows for long term preservation and storage of samples while retaining useful morphological information. The process of fixation, embedding, and extraction can damage RNA, and long-term storage of such samples can make the damage progressively worse, making gene expression analysis difficult (4, 13). However, due to the nature of DO hybridization and ligation chemistry and the short length of RNA sequence that is targeted by each DO set, TempO-Seq is highly resistant to this type of fragmentation, as well as to the presence of crosslinking.

To determine if storage time of FFPE blocks had a significant effect on performance of the FFPE TempO-Seq assay, we obtained archival human tumor FFPE samples from the University of Arizona Cancer Center Biorepository. Archival tissues, with their indicated year of harvest, were as follows: colorectal cancer (1986), hepatocellular carcinoma (1993), and two separate cases of kidney cancer (1994 and 1988). Blocks had been stored at room temperature, and in early 2016, were cut into 5µm thick sections. Slides were stored for two years before 25 mm^2^ areas were scraped for FFPE TempO-Seq processing using the human whole transcriptome panel. The same area was cut from serial sections to produce biological replicates. On average, each sample generated 2.1 M mapped reads, which is sufficient for meaningful data analysis (only a 50 base pair region is sequenced and counted for each gene using TempO-Seq, compared to RNAseq in which identification of each gene requires sequencing and counting multiple fragments). We compared gene expression data between biological replicates as a read out of assay reproducibility. Each of the archival samples had R^2^ values of greater than 0.8, with the Kidney harvested in 1994 having a biological replicate R^2^ of 0.925 (Figure 4). This finding demonstrated that the FFPE TempO-Seq assay can produce robust data sets from FFPE samples that are more than 30 years old.

**Figure 4.**
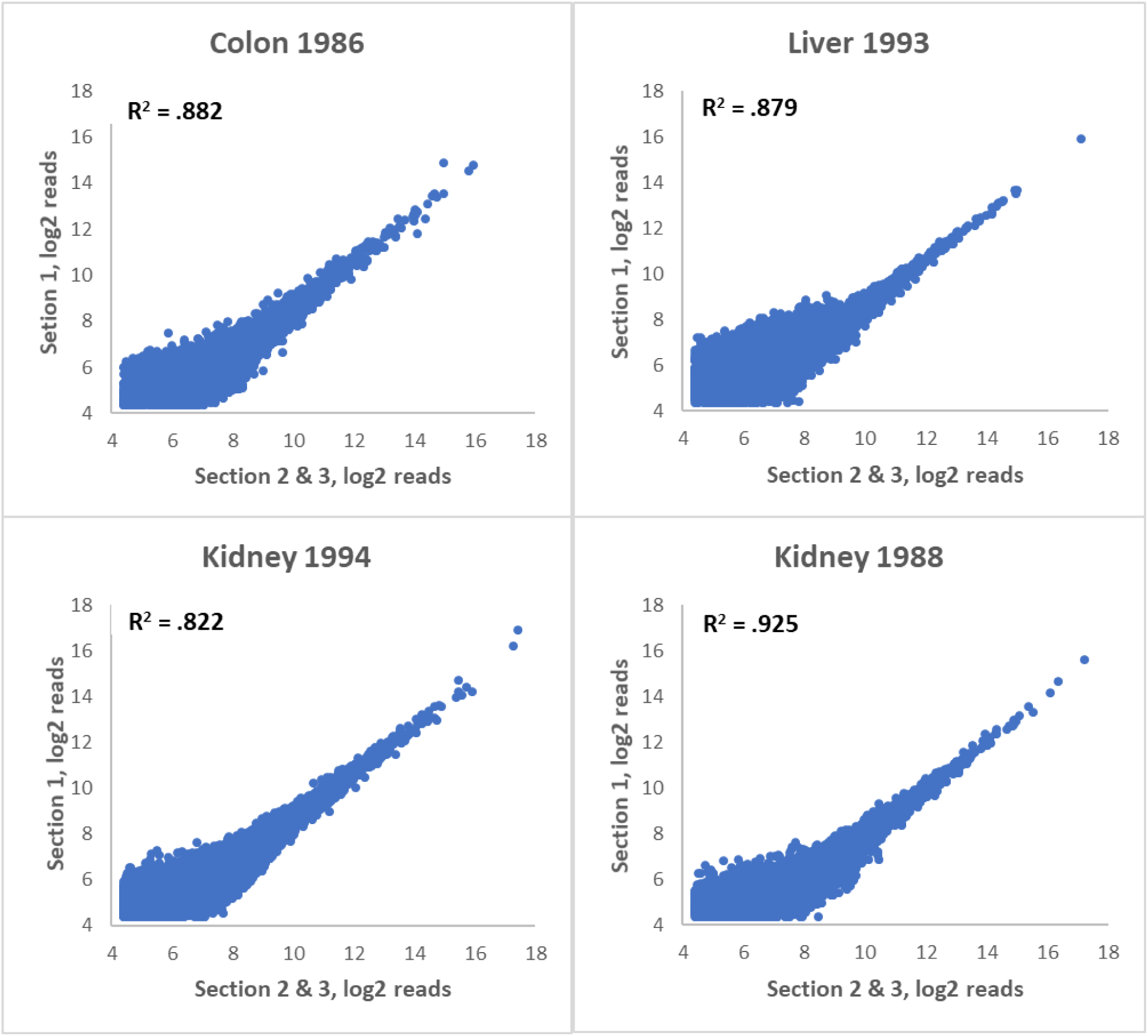
Gene expression correlation in biological replicates of archived human FFPE tissue. Human tissues were harvested in the year indicated and stored as FFPE blocks. Tissue sections were scraped and input into the FFPE TempO-Seq Human Whole Transcriptome assay. Replicates were obtained by scraping adjacent areas from three different serial sections. R^2^ values were calculated by comparing one section to the average of the other two.

### Fixation time

The amount of time tissue is exposed to fixative correlates with tissue autolysis and damage caused by endogenous endonucleases. Furthermore, total time of fixation affects RNA integrity, which directly impacts cDNA synthesis from RNA derived from fixed tissues (3, 4). This factor can significantly confound gene expression analysis which relies on methods dependent on reverse transcription such as microarrays or RT-PCR. Furthermore, additional fixation time can lead to overfixation, affecting accessibility of RNA (4, 14, 15). We tested whether fixation time had a notable impact on our assay by harvesting rat liver tissue, incubating in 10% neutral buffered formalin at 4°C for 24, 96, 192, and 384 hours before embedding. 10mm^2^ of tissue was scraped from 5µm thick sections and used as input for the FFPE TempO-Seq assay using rat whole transcriptome DOs.

There was no negative effect on sequencing quality with additional fixation time beyond 24 hours. Gene expression between biological replicates was high: R^2^ = 0.96 for 24 hours; 0.93 for 96 hours; 0.95 for 192 hours, and 0.98 for 384 hours (Figure 5). For all fixation times, the observed expression pattern clearly matched that expected for hepatocytes. These data collectively demonstrate that the TempO-Seq assay performs robustly even on samples that have been fixed for extended periods of time.

**Figure 5.**
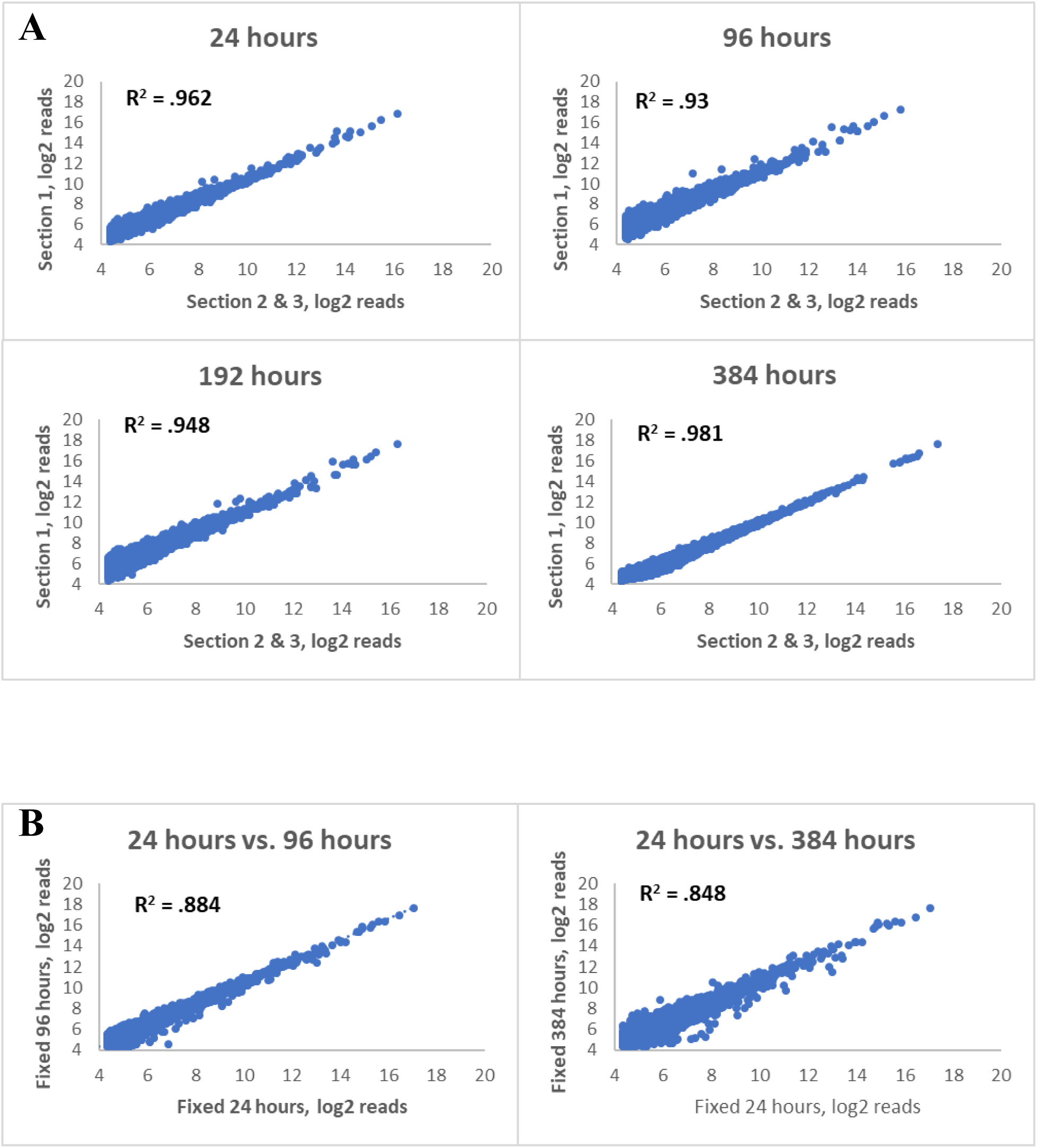
Gene expression correlation in biological replicates of rat FFPE tissues fixed for variable amounts of time. (**A**) Rat liver tissue was harvested and fixed in 10% neutral buffered formalin for 24, 96, 192, and 384 hours. Samples were then moved into 70% ethanol and processed for FFPE and used as input into the FFPE TempO-Seq rat whole transcriptome assay. Replicates were obtained by scraping adjacent areas from three different serial sections. R^2^ values were calculated by comparing one section to the average of the remaining two. (**B**) Comparison between 24-hour fixation and 96 or 384 hour fixation. R^2^ values were calculated between averages of three biological replicates for each condition.

### FFPE samples vs. fresh samples

Since fixation denatures RNA-binding proteins and disrupts secondary structure, TempO-Seq probes may interact with RNA in the context of fixed tissue differently than in fresh tissue. We compared the FFPE TempO-Seq assay to the standard assay designed for fresh lysates or purified RNA to determine whether processing of FFPE samples may impact biological conclusions. FFPE cell pellets were made from MCF-7 and MDA-MDA-231 breast cancer cell lines, derived from luminal A and claudin low subtypes, respectively. 10mm^2^ areas were excised from 5 µm thick sections and used as input into the FFPE assay. Cells from the same plate were lysed fresh in lysis buffer and used as input into the standard TempO-Seq assay (7).

We conducted differential gene expression analysis using DESeq2 between the two cell types for both FFPE and fresh assays. Differentially expressed genes were defined as genes with raw counts > 20, and by p_adj_ < 0.05. A total of 4,461 genes were detected as differentially expressed in FFPE samples using these cutoffs, compared to 3,015 in fresh lysates.

The log2 fold change between the two sample types showed a strong correlation (R^2^ = 0.970), (Figure 6A). Literature and previous gene expression data for these two cell types agree with genes detected by TempO-Seq as differentially expressed in both sample types (7).

**Figure 6.**
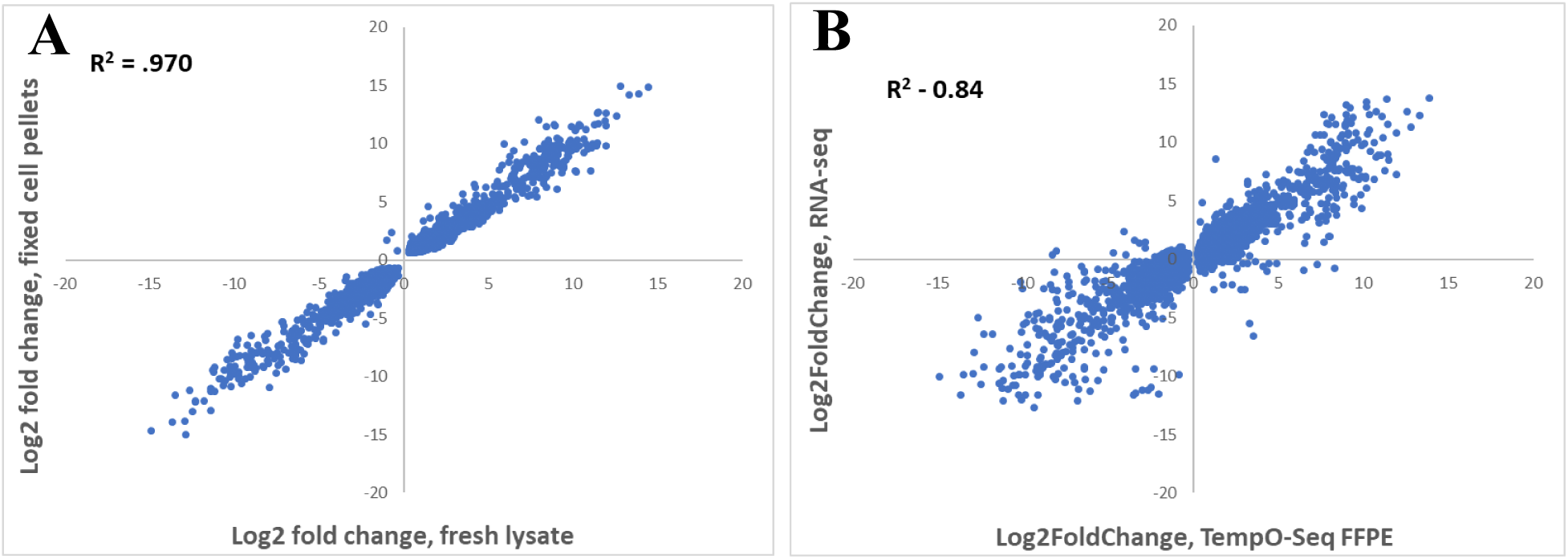
Correlation of gene log2 fold changes between fresh and fixed MCF-7 and MDA-MDA-MB-231 cells, and correlation of TempO-Seq with RNA-seq. (**A**) Both MCF-7 and MDA-MB-231 cells from the same plate were split in half. One half was lysed directly and processed through the standard TempO-Seq human whole transcriptome assay (fresh lysate), and the other half was pelleted and fixed for FFPE embedding and sectioning to be processed through the FFPE TempO-Seq human whole transcriptome assay (fixed cell pellet). For each assay, differential expression was calculated between the two cell types. Log2 fold changes were then plotted for each assay to determine correlation of differential expression. (**B**) RNA-seq was performed on RNA purified from fresh MCF-7 and MDA-MB-231 cells. FFPE TempO-Seq assay was performed on FFPE cell pellets produced from the same cell types. DeSeq2 was used to determine statistically significant (p_adj_ <0.05) log2 fold changes between the different cell types and plotted to derive correlation of differential gene expression between the two methods.

### Comparison to RNA-seq

While these data show that the TempO-Seq direct lysis assay of FFPE is reproducible, precise, and sensitive, the question of accuracy remains: how well do these measures reflect biological reality? RNA-seq is a method which depends on purification and reverse-transcription of RNA (both of which can introduce artifacts), and thus isn’t a perfect measure of biological reality. However, it has long been the gold standard for measurement of gene expression changes, and thus represents a sufficiently valid baseline measure for comparison.

We compared our MCF7 vs. MDA-MB-231 FFPE cell pellet results with previously published RNA-seq data (7) which measured log2FoldChange differences between RNA purified from the same cell types. We compared data points for genes with >20 counts, and whose expression was determined to be significantly different by DeSeq2 (p_adj_ <0.05). The agreement of log2FoldChange measures is excellent (R^2^=0.84), especially when differences in platforms and sample (RNAseq of extracted RNA from unfixed cells vs TempO-Seq assay of FFPE lysates) are taken into consideration (Figure 6B).

### Stained tissue

Hematoxylin and eosin (H&E) staining of FFPE sections is a practice commonly used for histopathological interpretation of tissue samples and can be used to identify a wide variety of diagnostically relevant features including cellular organization, nuclear morphology, and lymphocytic invasion (16). The H&E staining process requires deparaffinization and rehydration of tissue slides before staining. Samples that are rehydrated risk hydrolysis of RNA molecules and exposure to RNases, in addition to RNA degradation can occur due to relatively high acidity of the staining process. Therefore, current practice is to prepare an H&E stained section and then process an adjacent unstained section using the H&E section as a guide. This is fine so long as there is sufficient material and the whole slide is being processed. However, if only a focal area is of interest because of its histology within the section, then marking slides accurately based on a serial H&E stained section can be problematic, particularly if the area is very small. Therefore, we pursued the possibility of profiling the H&E stained slide itself, so that the area scraped could be directly visualized and documented.

To test whether H&E staining would interfere with TempO-Seq, we used a set of 5 µm thick serial human prostate cancer slides. These slides were either processed directly, deparaffinized and processed, or deparaffinized then H&E stained and processed. RNase-free reagents were used for deparaffinization and staining. The same 5 mm^2^ area of homogenous tumor was scraped from each serial section and lysed, with 2 mm^2^ equivalent of the resulting lysate used as input into the human whole transcriptome FFPE TempO-Seq assay. Gene expression profiles between paraffinized and deparaffinized sections had a high correlation (Fig 7, left panel, R^2^ = 0.902), indicating that the method of paraffin removal had little effect on assay performance. The R^2^ value between deparaffinized and H&E stained was also high (R^2^ = 0.855), demonstrating that the assay still works well with RNA exposed to the H&E chemistry. Gene expression signatures from H&E stained tissue also correlated well with unstained sections (R^2^ = 0.841). Overall, these data demonstrate that H&E stained FFPE tissue can serve as input for the TempO-Seq assay as long as RNase free reagents are used in the H&E staining process. This also further validates the ability of the assay to detect significantly degraded RNA within samples (Figure 7).

**Figure 7.**
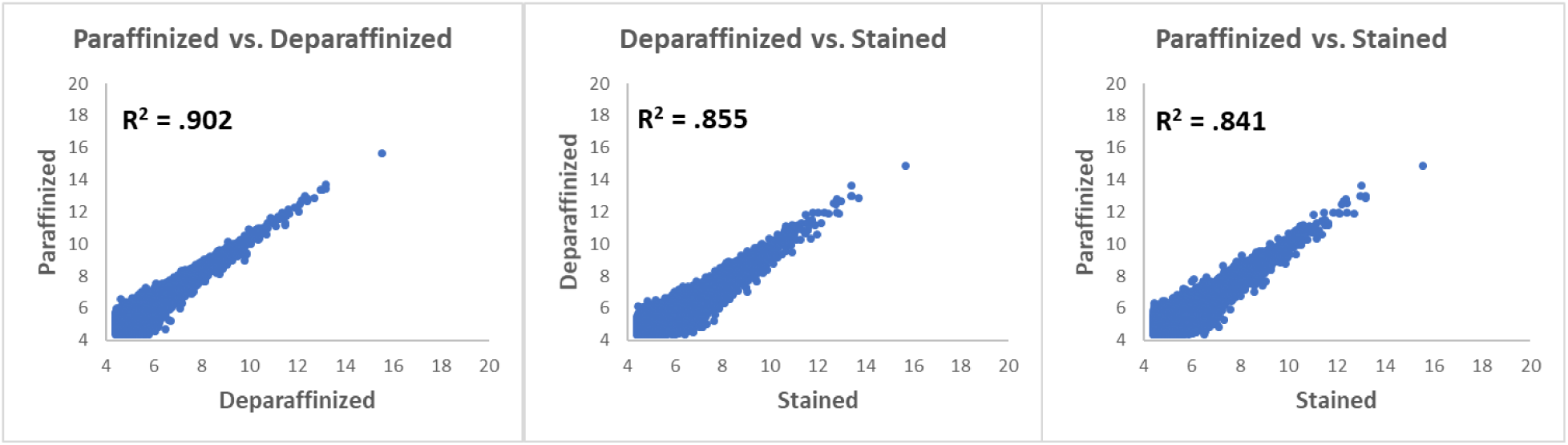
Gene expression correlation of human prostate FFPE samples that were unstained, deparaffinazed, or deparaffinized and H&E stained. Serial sections were scraped and used as input into the FFPE TempO-Seq assay for human whole transcriptome. Gene expression correlation was conducted to compare paraffinized (sectioned and mounted, but not otherwise treated), deparaffinized, and deparaffinized then stained conditions.

## Discussion

The processing required for fixation and paraffin embedding of tissues tends to fragment nucleic acids, which presents significant obstacles to molecular analysis. This is particularly the case for RNA and measurement of gene expression. However, FFPE samples also preserve tissue morphology well over long periods of time, and are easy to handle and section, factors that make them extremely useful for pathology and long-term storage. Additionally, after the initial damage induced by the process itself, fixation and embedding provide significant protection from further damage and hydrolysis, without the need for expensive or cumbersome measures such as snap-freezing and keeping samples constantly frozen for years or decades. These advantages have led to accumulation of vast archives of annotated FFPE tissues which have until now been difficult or impossible to profile at the molecular level.

TempO-Seq (4, 7-11) is a targeted, ligation-based assay designed to minimize complexities usually associated with gene expression measurements. Since it processes and counts only specific pre-determined probe sequences, it avoids cumbersome bioinformatics (the output of the assay is a simple table of counts for each gene in each sample) and reduces sequencing costs significantly (to 1/10^th^ or less). It can be performed in any lab, requiring no specialized equipment beyond a thermocycler and access to a sequencing instrument (commercially, or in most university core facilities). By avoiding RNA extraction and reverse transcription, the processing of samples is simplified, requires less hands-on time, and is relatively insensitive to fragmentation because only a <100 nt sequence of the RNA is targeted. All these factors allow TempO-Seq to directly measure gene expression in a wide variety of FFPE samples, and from small amounts of FFPE.

In this study, we provide data demonstrating the quality and reproducibility (Figure 2, Table 1) of the TempO-Seq assay of a variety of FFPE tissue samples across three different species (human, mouse, and rat). The gene expression data agrees well with existing data from the literature (Table 2), and is highly sensitive, producing excellent gene expression readouts from tissue inputs as small as 2 mm^2^ areas of a 5 µm section (Figure 3). In comparison, most other methods require sacrifice of entire sections (or multiple sections) to obtain sufficient extracted RNA for a single attempt at measurement. This level of sensitivity is critical when dealing with precious and irreplaceable archival samples. The assay is also insensitive to the time samples are stored (Figure 4) or the length of fixation (Figure 5).

While it is important to demonstrate the performance within FFPE and between different tissues, it is just as important that the assay of FFPE provide the same results as from unfixed tissue. This was demonstrated in Figure 6A, where the data for unfixed and fixed showed an excellent correlation (R^2^ = 0.97). These experiments demonstrate that the FFPE data is reliable and as accurate and meaningful as data from profiling unfixed tissue. This is true not only within the TempO-Seq platform, but as Figure 6B demonstrated, between differential expression measured using RNAseq and the TempO-Seq platform, producing a “between platform” R^2^ = 0.84. This is notable, particularly considering that typical between-platform correlations assay the same sample (e.g. aliquots of same extracted RNA); while in this case we compared the RNA-seq differential expression of RNA extracted from unfixed cells to the whole transcriptome TempO-Seq assay of cell pellets after they were fixed, embedded in paraffin, and then sectioned before assay. This combined cross-platform and cross-methodology consistency demonstrates that results from this assay are likely to reflect the true expression profile of any assayed sample.

The observed sensitivity to assay small areas of FFPE (Figure 3) becomes especially important when coupled with data showing that TempO-Seq can be performed on H&E stained tissues (Figure 1A and 7), as long as the staining is performed using RNase-free reagents. In practice, this means individual tissue sections can be stained, and the staining used to determine precisely delimited areas of tissue to be profiled (e.g. separating epithelial cells from background, or stromal tissue from glands, etc.). Examples of H&E stained tissue are shown in Figure 1A, where the heterogeneity of the FFPE is evident (upper right panel) as well as the consistency of histology within the small scraped area (lower left and right panels). The ability to profile small focal areas of H&E stained FFPE, targeting specific histological features within the section to obtain whole transcriptome data is unique to TempO-Seq. It should enable investigators to obtain highly histology-specific gene expression data to delineate not only disease states but also the complex interactions between cell types and histologies within a tissue.

The combined sensitivity, robustness, and consistency of expression profiling permit the TempO-seq FFPE whole transcriptome assay to be used over a wide variety of applications which would not otherwise be possible. By opening the doors to many previously inaccessible samples and study designs, we believe that use of the whole transcriptome TempO-Seq assay of FFPE samples will lead to significant advancements in many fields of biological science.

## Acknowledgments

Funding for the development of the FFPE TempO-Seq assay was made possible by National Institute of Health National Cancer Institute grants 5R33CA183688-02, and National Institute of Environmental Health Sciences grant 1R43ES024107-01. The authors would like to thank Ditte Andersen and Euan Cameron from BioClavis, Inc. (UK) for insight into assay development and experimental design. We thank Dr. Ray Nagel from the University of Arizona Department of Pathology for identifying tumor regions in archival samples. We also thank Kathleen Scully and Pamela Itkin-Ansari of the Sanford Burnham Prebys Medical Discovery Institute for providing mouse tissues. Finally, we thank Marilyn Maron, Marisa Gonzalez, and Khue Tran from BioSpyder Technologies for providing reagents, feedback, and resources for data analysis.

